# Fragile X Mental Retardation Protein (FMRP) expression in human nociceptor axons of the spinal dorsal horn— Implications for RNA targeting and localized translation

**DOI:** 10.1101/2022.09.15.508178

**Authors:** Molly E. Mitchell, Lauren C. Cook, Stephanie I. Shiers, Diana Tavares-Ferreira, Armen N Akopian, Gregory Dussor, Theodore J Price

## Abstract

Fragile X Mental Retardation Protein (FMRP) regulates activity-dependent RNA localization and local translation to modulate synaptic plasticity throughout the CNS. Mutations in the *FMR1* gene that hinder or ablate FMRP function cause Fragile X Syndrome (FXS), a disorder associated with sensory processing dysfunction. FXS pre-mutations are associated with increased FMRP expression and neurological impairments including sex dimorphic presentations of chronic pain. In mice, FMRP ablation causes dysregulated DRG neuron excitability and synaptic vesicle exocytosis, spinal circuit activity, and decreased translation-dependent nociceptive sensitization. Activity-dependent, local translation is a key mechanism for enhancing primary nociceptor excitability which promotes pain in animals and humans. These works indicate that FMRP likely regulates nociception and pain at the level of the primary nociceptor or spinal cord. Therefore, we sought to better understand FMRP expression in the human dorsal root ganglion (DRG) and spinal cord using immunostaining in organ donor tissues. We find that FMRP is highly expressed in DRG and spinal neuron subsets with substantia gelatinosa exhibiting the most abundant immunoreactivity in spinal synaptic fields. Here, it is expressed in nociceptor axons. FMRP puncta colocalized with Nav1.7 and TRPV1 receptor signals suggesting a pool of axoplasmic FMRP localizes to plasma membrane-associated loci in these branches. Interestingly, FMRP puncta exhibited notable colocalization with calcitonin gene-related peptide (CGRP) immunoreactivity selectively in female spinal cord. Our results support a regulatory role for FMRP in human nociceptor axons of the dorsal horn and implicate it in the sex dimorphic actions of CGRP signaling in nociceptive sensitization and chronic pain.

## 1 Introduction

For millions of individuals, chronic pain remains a devastating, daily challenge that dramatically reduces their quality of life (Dahlhamer, 2018; Yousuf et al., 2021). Notably, the magnitude and especially the frequency and duration of these symptoms are significantly magnified in women when compared to men (Fillingim et al., 2009; Mogil, 2020; Traub & Ji, 2013). With available pharmacological therapies demonstrating only modest efficacy at best, outcomes for chronic pain patients remain remarkably poor (Finnerup et al., 2015; Pagé et al., 2018; Siracusa et al., 2021; Yousuf et al., 2021). A major obstacle is that few findings from pre-clinical rodent models are successfully translated into clinical treatments (Abboud et al., 2021; Clark, 2016; Lenert et al., 2022). A core concern is our limited understanding of the cellular and sub-cellular organization of proteins expressed in human nociceptive circuitry that have critical regulatory roles in pain pathways in humans. Primary nociceptors in the DRG are the first cells of the pain pathway, and their activity is required for the generation of most pain symptoms in both acute and chronic pain (Haroutounian et al., 2014, 2018; Khoutorsky & Price, 2018; Price & Géranton, 2009; Price & Gold, 2018; Siracusa et al., 2021; Yousuf et al., 2021). Damage-associated signals change the excitability of nociceptors through a process that includes changes in gene expression at the level of both somatic and axonal mRNA translation (Khoutorsky & Price, 2018; Megat & Price, 2018; Price & Géranton, 2009; Reichling & Levine, 2009; Yousuf et al., 2021).

With critical functions in both activity-dependent somatic and local translation in neurons, Fragile X Mental Retardation Protein (FMRP) is likely an important regulator of changes in gene expression in nociceptors. *FMR1* mRNA is expressed in both male and female human nociceptors, and the protein is abundant in the somata and axons of rodent nociceptors (Price et al., 2006, 2007; Price & Melemedjian, 2012; Tavares-Ferreira, Shiers, et al., 2022). FMRP is an RNA-binding protein (RBP) that is indispensable for several forms of long-term plasticity, including nociceptive sensitization in rodent pain models (Asiedu et al., 2011; Johnson et al., 2022; Mei et al., 2020; Price et al., 2007; Ramírez-López et al., 2021; L. Yang et al., 2020; Y. Yang et al., 2021). Loss of function mutations in *FMR1* cause Fragile X Syndrome (FXS) (Collins et al., 2010; Darnell & Klann, 2013; De Boulle et al., 1993; Hagerman et al., 2017; Lozano et al., 2016; Myrick et al., 2015). People with FXS often exhibit severe sensory processing deficits including diminished nociceptive responses (Hagerman et al., 2017; Lozano et al., 2016; Symons et al., 2003). In line with this, *FMR1* null mice exhibit decreased nociceptive sensitization including in inflammatory and neuropathic pain models (Asiedu et al., 2011; Price et al., 2007).

FXS is the best-known disorder associated with *FMR1* mutations and mostly arises from expansion of the CGG trinucleotide repeat in its 5’UTR resulting in the transcriptional silencing of the gene and loss of FMRP (Jin & Warren, 2000; Peprah, 2012). Premutation is associated with fewer CGG repeats and increased FMRP expression (Jacquemont et al., 2004; Tassone et al., 2007). Premutation carriers can develop Fragile X-Associated Tremor Ataxia Syndrome (FXTAS) and frequently present with chronic pain (Au et al., 2013; Hagerman et al., 2007; Jalnapurkar et al., 2015; Johnson et al., 2022; Leehey et al., 2011; Salcedo-Arellano et al., 2020). In these populations, *FMR1* expression is increased (2- to 8-fold) likely due to enhanced transcription (Jacquemont et al., 2004; Storey et al., 2021; Tassone et al., 2007). FXTAS is associated with back pain and peripheral neuropathy with both being potential presenting features of the disorder (Hagerman et al., 2007; Johnson et al., 2022; Salcedo-Arellano et al., 2020). Premutation females exhibit notably high incidences of certain chronic pain disorders including fibromyalgia (43%) and migraine (F: 54.2%; M: 26.8%) (Au et al., 2013; Leehey et al., 2011). Collectively, these studies provide strong, bi-directional evidence for a role of FMRP in pain. Evidence from rodent models suggests that the likely site of action is the DRG or spinal cord (Asiedu et al., 2011; P.-Y. Deng et al., 2021; Ferron et al., 2020a; Mei et al., 2020; Price et al., 2006, 2007; Ramírez-López et al., 2021; Y. Yang et al., 2021). Therefore, we sought to better understand FMRP expression in the human DRG and spinal cord using tissues recovered from organ donors.

The objective of this study was to thoroughly characterize FMRP expression in human nociceptors from male and female organ donors with a focus on the spinal branches of their axons that project to the dorsal horn. We provide evidence that FMRP is present in human DRG somata and across the extent of these arbors. Our analyses also uncovered evidence of a sex dimorphism in the spinal presynaptic localization of FMRP to calcitonin gene-related peptide (CGRP) loci in the female spinal cord. Our results support the hypothesis that FMRP likely regulates RNA targeting and local translation in the central axons of nociceptors in humans, a mechanism that is likely differentially engaged across nociceptor populations with potential sex dimorphic roles.

## 2 Methods

### 2.1 Tissue Preparation

All human tissue recovery procedures were performed as approved by the Institutional Review Board at the University of Texas at Dallas. Human DRG and spinal cord were recovered from neurologic determination of death organ donors in collaboration with the Southwestern Transplant Alliance (STA). Donor information is provided in **Table 1**. Tissues were recovered within 4 h of cross-clamp and immediately flash frozen in powdered dry ice before storage at −80°C (Shiers et al., 2021). Fixation was performed in a manner emulating that carried out by brain banks for human cortex and hippocampus samples previously used for FMRP immunostaining (Akins et al., 2017; *SFARI | Autism BrainNet*, 2015). The only optimization was that STA procedures permitted all tissues to remain frozen until fixation. DRGs and spinal cord were warmed to −20°C before cryosectioning at 60 μm for free-floating cross sections. Sections were immediately immersed in ice cold 4% paraformaldehyde (PFA) (120 mM phosphate buffer, pH 7.4 with 150 mM saline) for 15 min. They were then rinsed three times with 1× PBS, pH 7.4 (10 mM phosphate; 150 mM NaCl) and stored in 1× TBS with 0.02% sodium azide at 4°C until immunostaining.

**Table 1.** Donor demographics

| Donor | Tissue | Age | Cause of Death |
| --- | --- | --- | --- |
| Male 0 | DRG<br>*Lumbar 4* | 37 | Cerebrovascular accident/Stroke |
| Male 1 | Spinal cord<br>*Lumbosacral* | 19 | Anoxia/Drug Overdose |
| Male 2 | Spinal cord<br>*Thoracolumbar* | 63 | Head Trauma/Blunt Injury |
| Male 3 | Spinal cord<br>*Lumbar* | 53 | Anoxia/Myocardial infarction |
| Male 4 | Spinal cord<br>*Thoracolumbar* | 19 | Head Trauma/GSW |
| Female 0 | DRG<br>*Lumbar 4* | 57 | Anoxia/Cardiac Arrest |
| Female 1 | Spinal cord<br>*Lumbosacral* | 61 | Anoxia/Cardiac Arrest |
| Female 2 | Spinal cord<br>*Thoracolumbar* | 44 | Anoxia/Cardiac Arrest |
| Female 3 | Spinal cord<br>*Lumbar* | 18 | Urea cycle metabolism disorder:<br>Anoxia/seizure; hepatic<br>encephalopathy |
| Female 4 | Spinal cord<br>*Thoracolumbar* | 16 | Head Trauma/MVA |

\*Note: Donor information is given for all samples that were used for all experiments.

### 2.2 Antibodies

For FMRP IF, the following antibody concentrations were used: 2F5-1 [1:500; aa 1-204 of human FMRP; DSHB 2F5-1 supernatant (Akins et al., 2017; Christie et al., 2009; Gabel, 2004)], 1C3 (1:500; FMRP N-terminus; Chemicon MAB2160), or 7G1-1 [1:1000; aa 354-368 of FMRP; DSHB concentrate (V. Brown et al., 1998, 2001; Christie et al., 2009; Price et al., 2006, 2007). All three monoclonal FMRP antibodies were previously validated for immunofluorescence in brain sections from *FMR1* null mice (Christie et al., 2009; Gabel, 2004). 7G1-1 was previously validated for immunofluorescence in DRG and spinal cord sections from *FMR1* null mice (Price et al., 2007). For comparisons of DRG and axonal FMRP IF across FMRP antibodies, 2F5-1, 1C3, or 7G1-1 was incubated with rabbit anti-TRPV1 (1:500; ThermoFisher PA1-748) or chicken anti-peripherin (1:1000; Encor CPCA-Peri), respectively (**Figure 3**). Dual-IF with nociceptor markers was performed by incubating 7G1-1 with either rabbit anti-TRPV1 or rabbit anti-CGRP (1:500; Immunostar 24112). For triple IF, mouse IgG_1_ anti-Nav1.7 (1:500; NeuroMab MABN41) was included. All secondary antibodies were raised in goat, purchased from ThermoFisher, and used at 1:200 (Akins et al., 2017; Gingrich et al., 2018). Alexa 647-conjugated secondaries were used for detection of FMRP: goat anti- mouse IgG_1_ (A21240) was used for 1C3 or goat anti- mouse IgG_2b_ (A21242) for 2F5-1 or 7G1-1. For dual-IF with DRG axons, these were paired with Alexa 488-conjugated secondaries: goat anti-chicken (A11039) or goat anti-rabbit (A11034). For triple IF, Nav1.7 primaries were detected with anti- mouse IgG_1_ Alexa 555 (A21127). Bleed through into the FMRP channel was ruled out by dual-IF using anti- mouse IgG_1_ Alexa 488 (A21121).

### 2.3 Immunofluorescence (IF)

Human DRG and spinal cord cross sections were rinsed thoroughly with PBS. FMRP IF with 2F5-1 was performed as done previously to detect axonal FMRP-RNPs in human brain tissues using confocal microscopy (Akins et al., 2017). IF procedures for 1C3 or 7G1-1 were evaluated and optimized based on comparisons to 2F5-1 immunostaining in human DRG and spinal cord tissues. For antigen retrieval of the 2F5-1 epitope, sections were first transferred to 10 mM sodium citrate, *p*H 6.0 and heated at 75°C for 30 min (Akins et al., 2017; Christie et al., 2009). All sections were treated with an RNP-stabilizing blocking solution [1% blocking reagent for nucleic acid hybridization (Sigma 11096176001) in PBST (PBS, *p*H 7.4; 0.3% Triton X-100)] for 1h (Akins et al., 2017; Chyung et al., 2018; Gingrich et al., 2018). Sections were then treated with primary antibodies in blocking solution and incubated overnight at room temperature. 7G1-1 incubations were two nights with axonal and/or nociceptor marker primaries added after the first night. All primary and secondary antibodies with applicable specifications are listed in **Table 2**. For secondary antibody detection, tissues were washed in PBST and then incubated in blocking solution containing the appropriate secondaries for 1 h at room temperature. Sections were washed in PBST and then with PBS containing DAPI (1:10,000). Tissue processing and mounting were performed as previously optimized by super resolution of FMRP-RNPs within axons in rodent brain sections for standard confocal microscopy (Akins et al., 2017; Gingrich et al., 2018). This reliably enhances image resolution to the attainable limit of a confocal microscope for subcellular imaging of punctate FMRP IF. Tissues were washed thoroughly with PBS to remove any remaining DAPI or detergent. Sections were then immediately submerged in mounting media (4% n-propyl gallate, 85% glycerol and 10 mM phosphate buffer, pH 7.4) for 10 min to dehydrate the tissue without drying. This formula was refractive index-matched to nervous tissue and Olympus silicone immersion oil (SIL300CS-30CC) to maximize resolving power of Alexa 647 emission using a scanning confocal microscope (Akins et al., 2017; Gingrich et al., 2018; Hicks et al., 2022; Zheng et al., 2011). To ensure consistent reduction in working distance, sections were directly transferred onto coverslips (Globe Scientific Inc.; No. 1.5), allowed to settle, and excess mounting media was wicked away. Uncharged Super Frost slides with a small volume of mounting media (~100 μL) to preserve fluorescence were plated onto the coverslips and sealed with clear nail polish.

**Table 2.** List of antibodies and specifications used for immunofluorescence

| ***FMRP Immunoreactivity*** |  |  |  |  |  |  |
| --- | --- | --- | --- | --- | --- | --- |
| **Clone and/or Antibody** | **Target Epitope** | **Vendor** | **Catalog #** | **RRID** | **Specifications** | **IF Confirmed in Human Nervous Tissues** |
| 1C3<br>(Mouse IgG1) | N-terminus | Chemicon | MAB2160 | AB_2283007 | N/A | Unclear<br>(Present study: 2F5-1 Comparisons) |
| 2F5-1<br>(Mouse IgG2b) | N-terminus<br>(aa1-204) | DSHB<br>(Tarakoff Lab/Fallon Lab) | 2F5-1 Supernatant | AB_10805421 | 1:500;<br>Antigen retrieval | Cortex and Hippocampus<br>(Akins et al., 2017) |
| 7G1-1<br>(Mouse IgG2b) | KH-Domain<br>(aa354-368) | DSHB<br>(Warren Lab) | 7G1-1 Concentrate | AB_528251 | 1:1000;<br>Two-night incubation | DRG and Spinal Cord<br>(Present study: 2F5-1 Comparisons) |
| Goat anti-mouse IgG1 647 | 1C3 Fc | ThermoFisher | A21240 | AB_2535809 | N/A | N/A |
| Goat anti-mouse IgG2b 647 | 2F5-1 Fc;<br>7G1-1 Fc | ThermoFisher | A21242 | AB_2535811 | N/A | N/A |
| ***Axon and Nociceptor Marker Immunoreactivity*** |  |  |  |  |  |  |
| **Antibody** | **Vendor** | **Catalog #** | **RRID** | **Dilution** | **Confirmed IF in Human Nervous Tissues** |  |
| Chicken anti-peripherin | Encor | CPCA-Peri | AB_228443 | 1:1000 | DRG and Spinal Cord<br>(Shiers et al., 2020) |  |
| Mouse IgG1 anti-Nav1.7 (N68/6) | NeuroMab | MABN41 | AB_2184355 | 1:500 |  |  |
| Rabbit anti-TRPV1 | ThermoFisher | PA1-748 | AB_2209010 | 1:500 |  |  |
| Rabbit anti-CGRP | ImmunoStar | 24112 | AB_572217 | 1:1000 |  |  |
| Goat anti-chicken 488 | ThermoFisher | A11039 | AB_228443 | 1:200 |  |  |
| Goat anti-mouse IgG1 488 | ThermoFisher | A21121 | AB_2535764 | 1:200 |  |  |
| Goat anti-mouse IgG1 555 | ThermoFisher | A21127 | AB_2535769 | 1:200 |  |  |
| Goat anti-rabbit 488 | ThermoFisher | A11034 | AB_2576217 | 1:200 |  |  |

### 2.4 Confocal Imaging and Lipofuscin Identification

Sections were imaged on either an Olympus FV1200 or an Olympus FV3000RS confocal microscope. Lipofuscin is a highly autofluorescent structure inherent to human nervous tissues that permeates all channels with strongest emission in yellow to orange spectra range (Bandyopadhyay et al., 2014; Katz & Robison, 2002; Shiers et al., 2021; Yung et al., 2016). To avoid false positives, autofluorescence including lipofuscin was detected in an empty red channel with a 561 laser (auto-561) (**Figures 1-3**). Lipofuscin was then identified as auto-561 signal that colocalized with FMRP signal and was omitted from all analyses. Acquisition parameters for FMRP IF were set as done before to image axonal FMRP-RNPs using 2F5-1 in human brain and 1C3 or 7G1-1 in rodent tissues using the HiLo look up table (Akins et al., 2017; Christie et al., 2009; Chyung et al., 2018; Gingrich et al., 2018). Laser power and sensitivity (HV) were adjusted until the maximum signal intensity for each FMRP antibody condition was limited to only a few saturated pixels (12 Bit). Offset was set using 2F5-1 signal and was adjusted until blue pixels were present in the FOV. Gain was kept at the default setting of 1 for all images. Settings: 10×, 20×, and 40× (laser power ≤ 5%, HV ≤ 600, Gain = 1, offset = 5); 1.25× (laser power ≤ 20%, HV ≤ 650, Gain = 1, offset = 10). Observations noted at 1.25× were originally observed with higher power objectives. Autofluorescence settings were never below FMRP IF parameters. Acquisition parameters for all axon and/or nociceptor markers were set using the same approach that for FMRP IF and were within those previously used for imaging of human DRG and spinal cord with this Olympus FV3000RS (Shiers et al., 2021).

**Figure 1.**
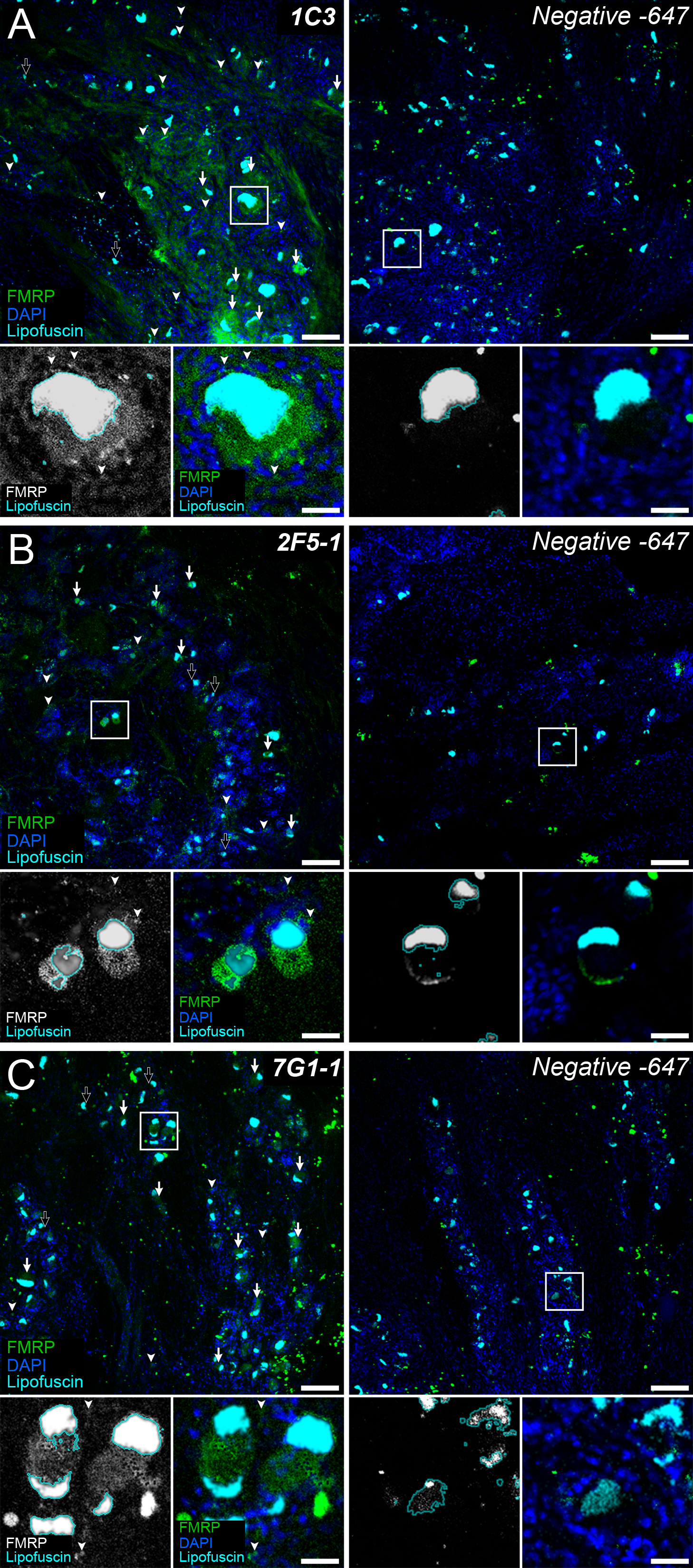
FMRP immunofluorescence in adult human dorsal root ganglion. Representative confocal micrographs showing FMRP immunofluorescence (**Green**) in adult human dorsal root ganglion using the 1C3 (**A**), 2F5-1 (**B**), or 7G1-1 (**C**) antibodies. DAPI counterstain (**Blue**) shows tissue organization. Lipofuscin were identified as outlined in methods (**Cyan**) and are present in all images. Negative controls were treated exactly as the matched FMRP immunostained tissue but were exposed only to Alexa 647-conjugated secondaries. For all FMRP primary antibodies tested, most DRG neuron somata exhibited FMRP IF (**Arrows**) with only a few appearing negative for FMRP (**Black Arrows**). These antibodies also exhibited signal consistent with FMRP staining in a subset of glia (**Arrowheads**). High magnification insets (**White Boxes**) show FMRP signal (**Insets: White or Green**) in DRG somata with surrounding glial nuclei (**Blue**). **A**: Robust 1C3 signal was detected in a large subset of neuronal somata and frequently in glial cells. **B**: Strong 2F5-1 signal was observed in most neuronal cell bodies and in some glia. **C**: Strong 7G1-1 signal was evident in neuronal somata and in few glia. Scale bars: Outsets = 100 μm; Insets = 30 μm.

### 2.5 Imaging for FMRP IF Comparisons

For FMRP IF evaluation in human DRG, two sections per FMRP antibody were compared at 10× and 20× magnification for two donors. Three-color 10× images with DAPI, auto-561, and FMRP were acquired to document comparisons (**Figure 1**). For human spinal cord, three sections per FMRP antibody from the same donor were compared at 1.25×, 10×, 20×, and 40× magnification. Two-color 1.25× and 10× images with auto-561 and FMRP and three-color 40× images including peripherin were acquired to document comparisons (**Figures 2** **and** **3**). Laser power and HV parameters were set as described above for 1C3, 2F5-1, or 7G1-1 and were used to compare maximum Alexa 647 signal intensities across these FMRP antibodies per magnification. 1C3 staining exhibited the brightest Alexa 647 signal with 2F5-1 and 7G1-1 signals appearing notably dimmer at 1C3 parameters for all magnifications. Setting these parameters using 2F5-1 or 7G1-1 signal consistently resulted in markedly saturated 1C3 signal whereas 2F5-1 and 7G1-1 signal were almost identical at all magnifications. 2F5-1 had been used before to detect FMRP puncta in human hippocampus but requires antigen retrieval, a process that often disrupts axonal and synaptic architecture (Akins et al., 2017; Alelú-Paz et al., 2008; Gingrich et al., 2018; Gutierrez-Mecinas et al., 2016; Jiao et al., 1999; O’Hurley et al., 2014). Differences detected between 2F5-1 or 7G1-1 and 1C3 were consistent with prior works showing that 1C3 has comparable affinities for both FMRP and the Fragile X Related Protein 1 that shares > 70% N-terminal homology with FMRP (Bauchwitz, 2009; Khandjian et al., 1998). 7G1-1 recognizes an epitope on FMRP that is not present in its paralogues (Darnell et al., 2009; Kirkpatrick et al., 2001).

**Figure 2.**
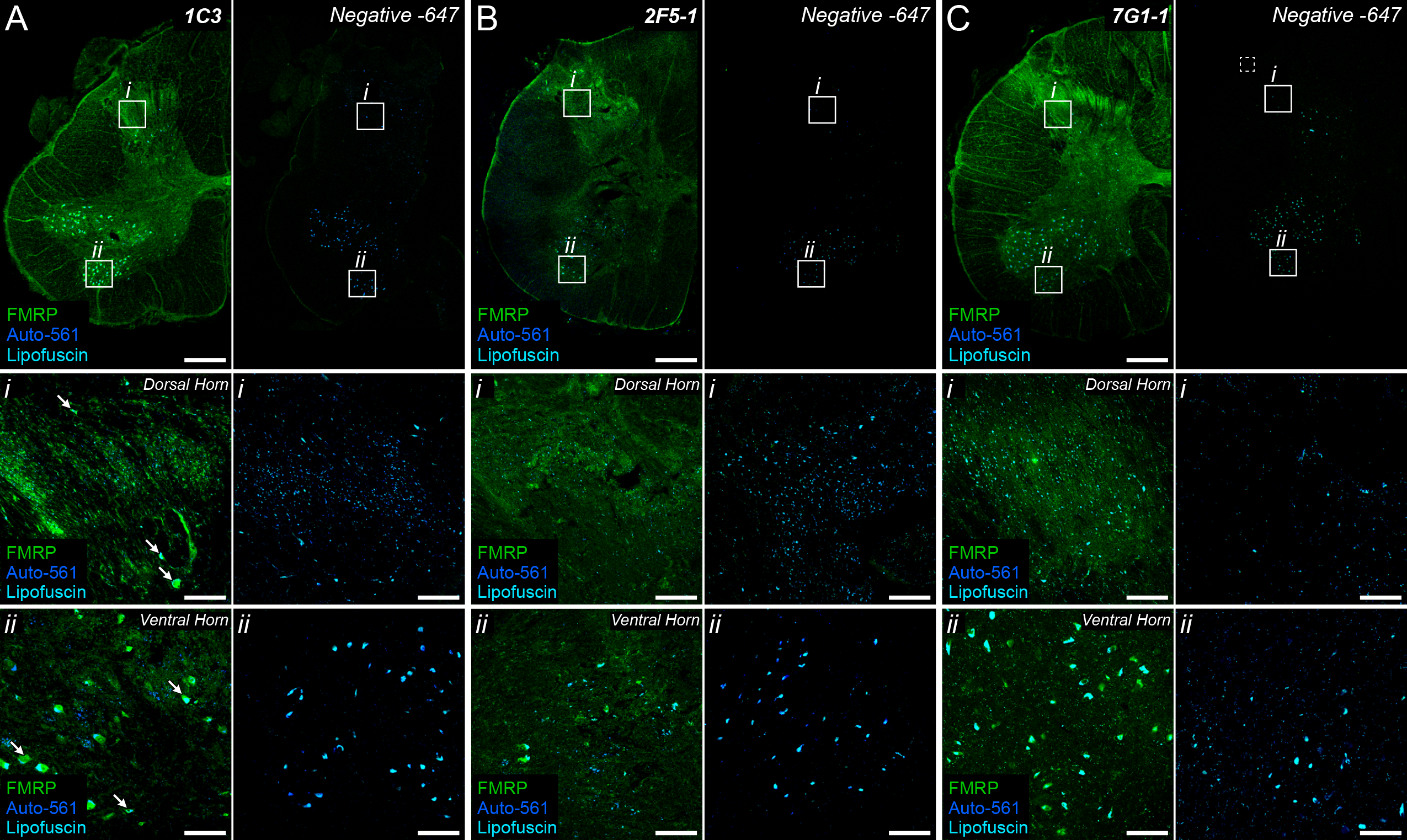
FMRP immunofluorescence in adult human spinal cord. Representative confocal micrographs of FMRP immunofluorescence (**Green**) in adult human spinal cord using the 1C3 (**A**), 2F5-1 (**B**), and 7G1-1 (**C**) antibodies. Autofluorescence (**Blue**) highlighted lipofuscin (**Cyan**). Negative controls were handled as described in Figure 1. 1.25× images of lumbar hemi-cross sections show grey matter regions with most intense FMRP signal per primary antibody. ***i-ii:*** 10× insets (**Boxes**) highlighting the consistency of increased FMRP signal in substantia gelatinosa (***Dorsal Horn***) or in lower motor neuron pools (***Ventral Horn***) across primary antibodies. Somatic signal was robust per antibody. **A**: 1C3 exhibited the brightest signal overall with the strongest signal in substantia gelatinosa (***i***) and motor neuron pools (***ii***). **B**: 2F5-1 signal was increased in neuropil within substantia gelatinosa (***i***) when compared to ventral grey (***ii***). Note the disrupted morphology due to antigen retrieval. **C**: 7G1-1 signal in somata and neuropil was comparable to that of 2F5-1 in substantia gelatinosa (***i***) and ventral grey (***ii***). Scale bars: Outsets = 1 mm; Insets = 100 μm.

**Figure 3.**
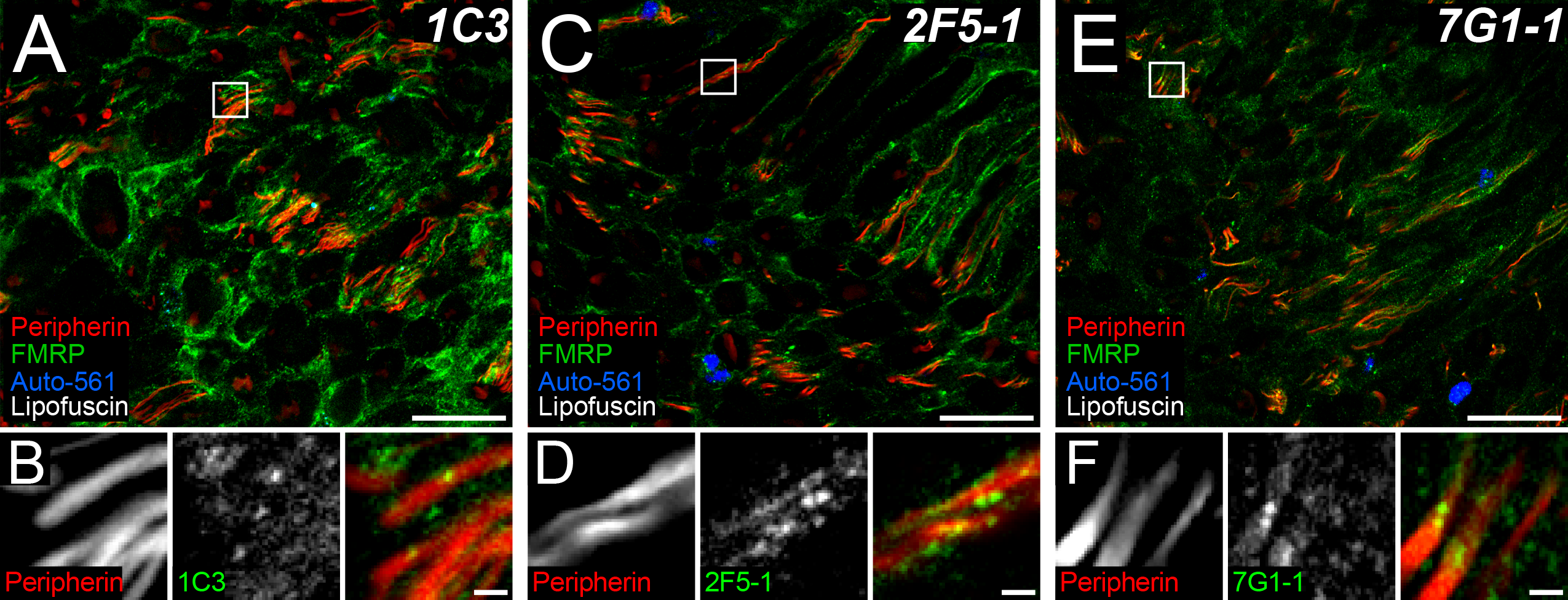
FMRP immunofluorescence in human dorsal root axons. Representative confocal micrographs of peripherin (**Red**) and FMRP (**Green**) immunostaining with 1C3 (**A**), 2F5-1 (**B**), or 7G1-1 (**C**) antibodies in human dorsal root. Autofluorescence (**Blue**) highlighted lipofuscin (**White**). High magnification insets (**Boxes**) show punctate FMRP signal colocalization with peripherin+ DRG axons for all FMRP antibodies tested. **A**: 1C3 signal was the most intense overall out the FMRP antibodies in dorsal root. **B**: The brightest 2F5-1 signal was often punctate as previously documented in axons within human hippocampus sections (Akins et al., 2017). **C**: 7G1-1 signal was similar to 2F5-1 signal. Scale bars: Magnified Images = 50 μm; Insets = 5 μm.

### 2.6 Imaging of FMRP IF and nociceptor marker colocalization

Four-color 40× images of dorsal root, lateral white matter tracts, and laminae I-II were acquired of spinal cord cross sections. To best avoid bias in our images, all imaging planes were set using the paired nociceptor marker channel by adjusting the focus to the focal plane exhibiting the most intense axonal signal that had the largest axon signal area to FOV area ratio. Maximum signal intensity across an FOV for each nociceptor marker was determined as described above for confocal imaging procedures. For all quantitative analyses, FMRP was detected with 7G1-1 and Alexa 647 as described above. For FMRP, peripherin, and Nav1.7 analyses, at least three images of dorsal root were acquired for each section, and three sections were imaged per donor for 2 donors (1 male and 1 female). Lipofuscin was identified by autofluorescence as described above but was detected with a 405 laser. Potential bleed through and differences in lipofuscin identification were ruled out by using dual-IF analysis of FMRP and Nav1.7 with auto-561 in two images from one section per donor. Images were acquired from 4 donors (2 male and 2 female) for FMRP and TRPV1 or CGRP dual-IF analyses with lipofuscin identified using auto-561. For TRPV1 in dorsal root or lateral tracts, one to two images per ROI were acquired from each section for three to four sections per donor. For CGRP analyses in dorsal root, two images were acquired from each section for two sections per donor. For all laminae I-II analyses, one to two images were acquired per medial, central, or lateral regions for three to five sections per donor.

### 2.7 Quantification of FMRP puncta and nociceptor marker colocalization

40× confocal micrographs had an approximate z-depth of 1 μm (~3% of the postfixation section thickness). An algorithm that was previously used to detect axonal FMRP-containing RNPs in brain tissues was optimized to detect FMRP puncta in human spinal cord sections (Akins et al., 2012, 2017; Gingrich et al., 2018). The original algorithm detected puncta with an apparent size of ~600 nm in 40× confocal micrographs that were ~200 nm by 3D-structured illumination super resolution microscopy (Akins et al., 2012, 2017). FMRP puncta detection in human spinal cord was restricted to those with a two-dimensional area of no less than 1 μm^2^. All quantification was performed in original micrographs. 12 Bit images were imported into FIJI (NIH, Bethesda, MD) and analyzed using the Analyze Particles function [1.00 – Infinity (microns); 0.00 – 1.00 (circularity)]. For each set of replicates and/or comparisons, a threshold of 205, 410, or 615 to 4095 set was used to detect 647 puncta in the FMRP channel FOVs including those that colocalized with nociceptor marker signal (488 or 555). The baseline autofluorescence for these cutoffs was based on the antigen retrieval-associated autofluorescence present in 2F5-1 treated spinal cord sections that did not meet FMRP puncta detection criteria in human brain tissues (Akins et al., 2017). The exact cutoff used was determined per experiment using the human tissue sample exhibiting the brightest baseline autofluorescence within an experimental set. This was done to best avoid quantification of 647 autofluorescence across replicates and/or comparisons given the intrinsic variability in sample quality. For detection of colocalized 647 puncta and nociceptor marker signal (488 or 555), puncta signal had a 100% overlap with nociceptor marker signal. Threshold cutoffs for marker signal were based on previous signal approximations and held constant across replicate and/or comparison sets (Shiers et al., 2020). All puncta varied greatly in size between donors and axonal compartments similar to that recently documented in axons within rodent hippocampus (Monday et al., 2022a). To generate an unbiased puncta count, puncta size was normalized to the total puncta area coverage within an FOV for each image for 647 and colocalized puncta. This generated a normalized count of the number of 1 μm^2^ puncta after dropping residual decimals which represent a partial punctum. Normalization examples using Male 3 dorsal root image 2:

647 puncta:

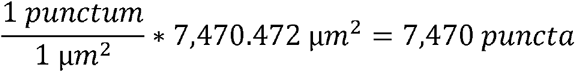

488-647 colocalized puncta:

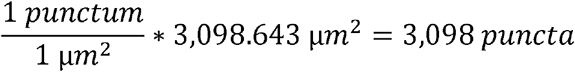

To avoid false positives due to lipofuscin, punctate lipofuscin signal was identified in the FMRP channel (647) or nociceptor marker channel(s) (488 or 555) using autofluorescence (auto-561 or auto-405) as described above. For total FMRP puncta counts, colocalization between punctate auto-signal and 647 signal was quantified using the parameters described for 647 puncta. Punctate lipofuscin size was then normalized to the total lipofuscin area per FOV and then subtracted from the normalized puncta count to generate total FMRP puncta counts. For colocalized FMRP puncta counts, this was also done for punctate auto-signal that colocalized with 488 or 555 signal. Averages for all puncta counts are listed in **Table 3**. FMRP and nociceptive marker IF covered large swathes laminae I-II ROI FOVs. To mitigated false positives for colocalization that may arise by chance, this number was approximated using the principles of Bayes’ Theorem. All calculations were carried out using lipofuscin subtracted total FMRP puncta or nociceptor markers areas (μm^2^). Puncta and areas for each channel shared a maximum possible area equal to the shared FOV area (μm^2^). Examples using Male 2 lateral laminae I-II image 2:

**Table 3.** Lipofuscin Identification and Subtraction from total and colocalized FMRP puncta

| Figure | Circuit ROI | Donor | Nociceptor Marker | Total Puncta (#) |  |  | Total Colocalized Puncta (#) |  |  |
| --- | --- | --- | --- | --- | --- | --- | --- | --- | --- |
|  |  |  |  | 647-Puncta | Auto in 647 | FMRP | Colocalized Puncta | Auto in Marker | Colocalized FMRP |
| 5 | Dorsal Root | Male 2 | Nav1.7 | 41,288 | 145 | 41,143 | 13,731 | 241 | 13,490 |
|  |  |  | Peripherin | 41,288 | 145 | 41,143 | 11,421 | 75 | 11,346 |
|  |  | Female 1 | Nav1.7 | 4,909 | 138 | 4,771 | 1,684 | 115 | 1,569 |
|  |  |  | Peripherin | 4,909 | 138 | 4,771 | 1,230 | 95 | 1,135 |
| 6 | Dorsal Root | Male 2 | TRPV1 | 17,647 | 118 | 17,529 | 16,638 | 136 | 16,502 |
|  |  | Male 3 |  | 29,604 | 380 | 29,224 | 12,641 | 816 | 11,825 |
|  |  | Female 1 | TRPV1 | 8,738 | 37 | 8,701 | 4,919 | 0 | 4,919 |
|  |  | Female 2 |  | 20,317 | 145 | 20,172 | 5,972 | 27 | 5,945 |
|  | Lateral Tracts | Male 2 | TRPV1 | 76,835 | 5,506 | 71,329 | 52,916 | 2,357 | 50,559 |
|  |  | Male 3 |  | 9,374 | 430 | 8,944 | 4,978 | 136 | 4,842 |
|  |  | Female 1 | TRPV1 | 2,632 | 302 | 2,330 | 565 | 0 | 565 |
|  |  | Female 2 |  | 28,265 | 1,143 | 27,122 | 9,676 | 208 | 9,468 |
|  | Laminae I-II | Male 2 | TRPV1 | 229,810 | 11,967 | 217,843 | 150,729 | 15,179 | 135,550 |
|  |  | Male 3 |  | 597,268 | 55,619 | 541,649 | 395,242 | 46,309 | 348,933 |
|  |  | Female 1 | TRPV1 | 40,119 | 4,763 | 35,356 | 15,528 | 381 | 15,147 |
|  |  | Female 2 |  | 384,771 | 17,335 | 367,436 | 143,254 | 2,651 | 140,603 |
| 7 | Dorsal Root | Male 2 | CGRP | 829 | 237 | 592 | 459 | 391 | 68 |
|  |  | Male 3 |  | 738 | 95 | 643 | 226 | 226 | 0 |
|  |  | Male 4 |  | 8,070 | 41 | 8,029 | 1,998 | 51 | 1,947 |
|  |  | Female 1 | CGRP | 18,199 | 88 | 18,111 | 8,375 | 124 | 8,251 |
|  |  | Female 3 |  | 20,760 | 61 | 20,699 | 8,478 | 63 | 8,415 |
|  |  | Female 4 |  | 11,868 | 113 | 11,755 | 8,881 | 133 | 8,748 |
|  | Laminae I-II | Male 2 | CGRP | 46,911 | 9,021 | 37,890 | 23,192 | 9,864 | 13,328 |
|  |  | Male 3 |  | 35,838 | 7,995 | 27,843 | 12,907 | 10,461 | 2,446 |
|  |  | Male 4 |  | 99,891 | 36,108 | 63,783 | 37,811 | 25,138 | 12,673 |
|  |  | Female 1 | CGRP | 76,827 | 14,120 | 62,707 | 47,450 | 13,542 | 33,908 |
|  |  | Female 3 |  | 55,012 | 7,374 | 47,638 | 30,254 | 6,282 | 23,972 |
|  |  | Female 4 |  | 319,063 | 23,468 | 295,595 | 138,096 | 11,100 | 126,996 |

\*Note: Lipofuscin were detected in both the FMRP and nociceptor marker channels by quantifying colocalization with either 561- (Figures 6 and 7) or 405- (Figure 5) induced autofluorescence using the same algorithm parameters to detect FMRP puncta. This was used to estimate the number of puncta arising from lipofuscin in channels of interest per FOV. This number was then subtracted from the evident number of puncta present in each FOV to more accurately identify punctate FMRP immunoreactivity. The example puncta numbers given are the overall sums detected and identified per donor for each quantitative analysis.

By chance adjustments for colocalized FMRP puncta:

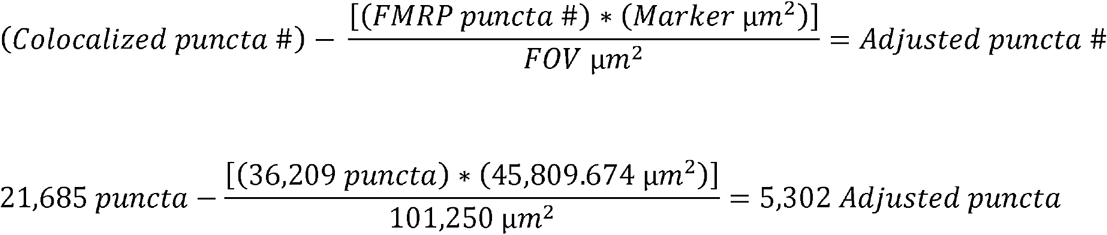

For detection of FMRP puncta and nociceptive marker colocalization in laminae I-II, colocalization using these criteria needed to be detected in dorsal root arbors.

### 2.8 Measurement of axonal diameter

40× images of peripherin, Nav1.7, and FMRP staining in dorsal root taken as described above were used to measure the approximate diameter of nociceptor axon arbors. Measurements were taken in the peripherin channel but were restricted to axonal sites exhibiting all three stains. This was done for all triple-IF sections and consisted of four to five axons per dorsal root image using both longitudinal and cross sections of axons. Four-color images were imported into FIJI and measurements were taken using the Line Tool function.

### 2.9 Proportion and Statistical Analysis

Pie charts and graphs were generated using GraphPad Prism version 9.4 (GraphPad Software, Inc. San Diego, CA USA). Male and female means for CGRP-colocalized FMRP puncta were compared using unpaired t-tests.

## 3 Results

### 3.1 FMRP immunoreactivity in the human dorsal root ganglion

We first sought to characterize FMRP immunoreactivity in human dorsal root ganglion (DRG). To do this, we analyzed FMRP immunofluorescence (IF) in lumbar DRGs from adult male and female organ donors (**Table 1**). We characterized FMRP IF with three different monoclonal antibodies generated against FMRP: 1C3, 2F5-1, or 7G1-1 (**Figure 1; Table 2**). Each of these antibodies binds to distinct epitopes on FMRP (**Table 2**) (Akins et al., 2017; V. Brown et al., 1998, 2001; Ceman et al., 2003; Christie et al., 2009; Devys et al., 1993; Gabel, 2004). For all antibodies, we found that many DRG neurons exhibited signal, even after accounting for lipofuscin (**Figure 1 A-C**) (Shiers et al., 2021). This was in line with prior observations in adult human hippocampal neurons and adult rodent DRG (Akins et al., 2017; Ferron et al., 2020a; Price et al., 2006, 2007). Some glial staining was evident for all FMRP antibodies, but it was most pronounced in 1C3-stained tissues (**Figure 1 A**). This consistent with observations that some glia express low levels of FMRP in postnatal rodent tissues (Gholizadeh et al., 2015; Pacey & Doering, 2007; Zorio et al., 2017). Overall, this indicates that FMRP is also localized to human DRG axons as previously observed in rodent tissues (Price et al., 2006).

### 3.2 FMRP immunoreactivity in the human spinal cord

We next determined whether FMRP is present in the human spinal cord. We immunostained for FMRP in adult human spinal cord sections exactly as we did in DRG (**Figure 2**; **Tables 1** **and** **2**). We found that each FMRP antibody exhibited IF signal in neuronal cell bodies present throughout the grey matter (**Figure 2 A-C**) (Akins et al., 2017; Christie et al., 2009; Kao et al., 2010; Monday et al., 2022b; Sawicka et al., 2019; Zimmer et al., 2017). IF in the superficial dorsal horn laminae appeared more abundant than that in both deep dorsal and ventral horn laminae (**Figure 2 A-C Hemi-sections and *i* vs *ii***). At high magnification, we also observed punctate staining in neuropil consistent with FMRP localization to pre- and post-synaptic processes (Akins et al., 2017; Christie et al., 2009; Kao et al., 2010; Monday et al., 2022b; Sawicka et al., 2019; Zimmer et al., 2017). These observations were comparable across the FMRP antibodies tested (**Table 2**). This indicates that in the dorsal horn, FMRP is present in neuronal somata but that synaptic FMRP is likely most abundant within the substantia gelatinosa. This region contains synapses that are comprised of connections between projection neurons and local interneurons as well as synapses of DRG afferents onto spinal cord neurons. To assess FMRP IF in DRG central axons entering the spinal cord, we examined sections of human dorsal root containing these arbors as identified by peripherin IF (**Figure 2 A-C; Table 2**) (Shiers et al., 2021). 1C3, 2F5-1, and 7G1-1 antibodies all exhibited strong and comparable signal in these DRG axonal compartments with 1C3 showing the strongest signal as in DRG. FMRP signal that colocalized with peripherin IF was predominantly punctate for the three FMRP antibodies (**Figure 3 A-C Boxes**). This was consistent with axonal FMRP-RNP staining in human and rodent brain tissues (Akins et al., 2017; Christie et al., 2009; Chyung et al., 2018; Gingrich et al., 2018; Monday et al., 2022b). This indicates that FMRP is present in the central branches of human DRG neurons and suggests that it is likely also present in the presynaptic structures of these axons in the spinal cord.

### 3.3 FMRP is present in DRG neuron afferent terminals in the dorsal horn

We immunostained spinal cord sections for FMRP (7G1-1) and peripherin, an axonal intermediate filament that is highly expressed by all peripheral neurons but not expressed by dorsal horn neurons (**Figure 4; Table 2**). We then examined colocalization between these two IF signals in confocal micrographs taken from different areas of the dorsal spinal cord (**Figure 4 A-F**). In the attached dorsal root, we found that punctate FMRP IF was present along the length of individual DRG axons (**Figure 4 B**). We also observed clusters of these puncta along peripherin+ axons in lateral white matter tracts (**Figure 4 C**). However, in the dorsal funiculus, we found that FMRP IF was virtually absent from DRG axons (**Figure 4 D**). This was in stark contrast to both laminae I and II where punctate FMRP staining was readily evident in neuropil, with lamina II exhibiting the strongest signal (Figure 4 E-F). FMRP IF also overlapped with peripherin+ spinal branches of DRG afferents in these laminae (**Figure 4 E-F Boxes**). In deep dorsal horn laminae V and VI-VII, we found that FMRP IF was present in cell bodies and at evidently lower levels in the associated neuropil (**Figure 4 G-H**). We detected puncta in a few peripherin+ DRG fibers in lamina V and rarely observed any in these arbors in laminae VI-VII (**Figure 4 G-H Boxes**). This suggests that in deep dorsal grey synaptic fields, FMRP IF likely arises in large part from immunoreactivity in dendritic compartments (Antar et al., 2004; Kao et al., 2010; Sawicka et al., 2019). Overall, our observations indicate that in the spinal branches of DRG neurons, FMRP IF is most abundant in those innervating laminae I-II.

**Figure 4.**
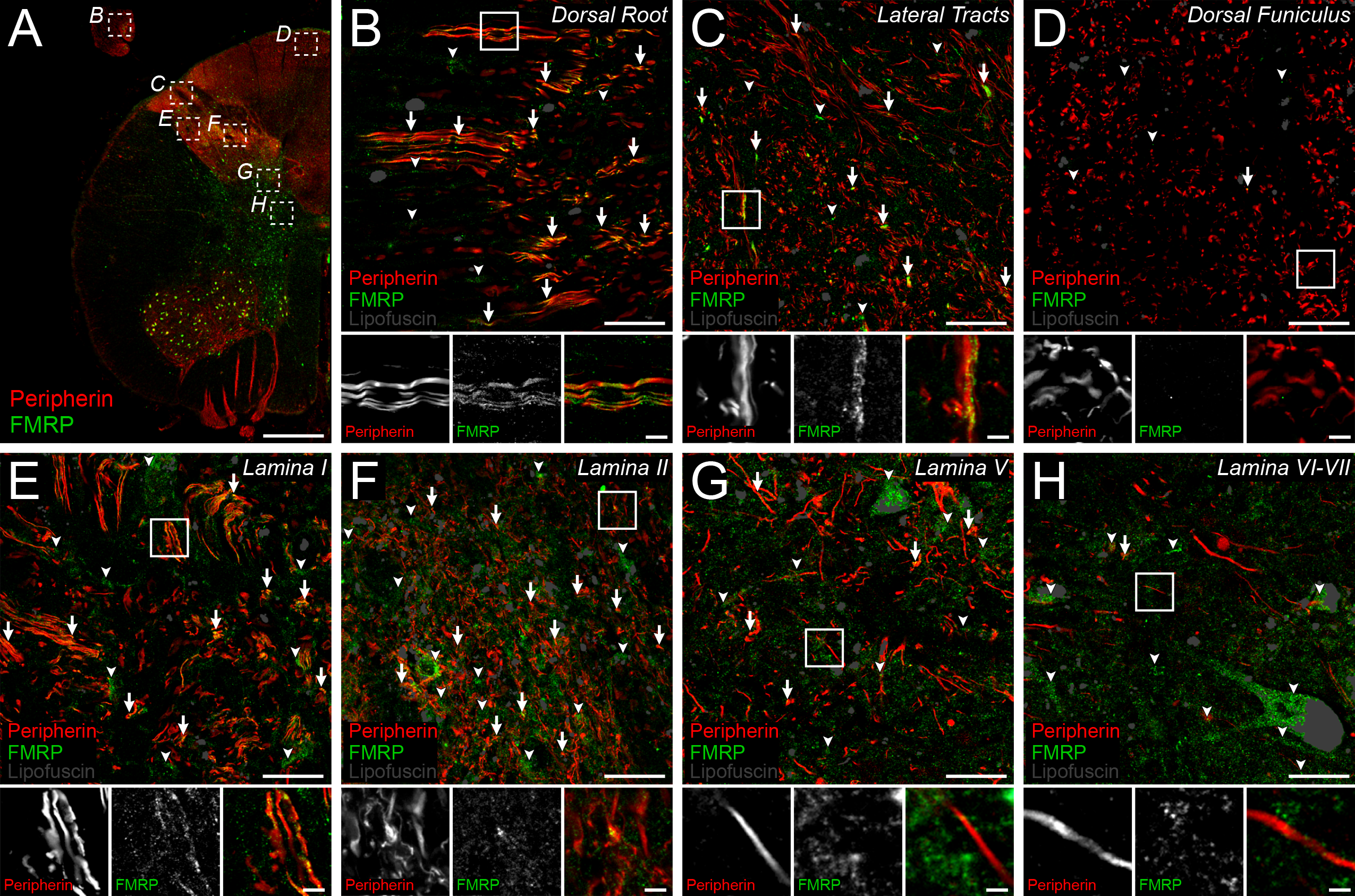
Peripherin and FMRP immunofluorescence in human dorsal horn. Representative confocal micrographs of immunostaining for peripherin (**Red**) and FMRP (**Green**) within dorsal horn in human spinal cord. Lipofuscin were identified as shown (**Grey**) to avoid false positives. **A**: 1.25× image of lumbar hemi-cross section depicting dorsal horn ROIs (**Dotted boxes B-H**) for 40× images used to assess peripherin and FMRP signal overlap. **B-H**: Magnified 40× images representing ROIs containing DRG axons and branches as identified by peripherin IF. The frequency of FMRP puncta and peripherin IF colocalization (**Arrows**) differed between ROIs. The proportion of FMRP puncta that did not colocalize with peripherin+ fibers (**Arrowheads**) was notably increased in grey matter ROIs. Associated insets (**Boxes**) are further magnified to highlight the sub-axonal morphology of signal overlap. **B**: Image of dorsal root shows FMRP puncta frequently colocalized with DRG axons. **C**: Image of lateral tracts show FMRP puncta often colocalized with peripherin+ axons at the spinal border of the DREZ. **D**: Image of dorsal funiculus shows virtually no colocalization between FMRP puncta and ascending peripherin+ arbors. **E**: Image of lamina II neuropil showing frequent colocalization between FMRP puncta and peripherin+ fibers. **F**: Image of lamina II neuropil shows FMRP puncta often colocalized with peripherin+ spinal branches in these synaptic fields. **G**: Image of lamina V neuropil shows dramatically reduced colocalization between FMRP puncta and peripherin IF. **H:** Image of lamina VI-VII neuropil shows FMRP puncta rarely colocalized with peripherin+ fibers in these synaptic fields. Scale bars: Outset = 1 mm; Magnified Images = 20 μm; Insets = 5 μm.

### 3.4 FMRP is present in plasma membrane microdomains in the spinal branches of primary nociceptors

Recent works have shown in brain that FMRP regulates axonal excitability and neurotransmission by directly binding and modulating ion channels in the axonal plasma membrane (M. R. Brown et al., 2010; P.-Y. Deng et al., 2013, 2019; P.-Y. Deng & Klyachko, 2021; Kshatri et al., 2020; Y. Zhang et al., 2012). In DRG neurons, it binds ion channels to regulate synaptic vesicle exocytosis and modulates their presence in the plasma membrane to regulate excitability (P.-Y. Deng et al., 2021; Ferron et al., 2014; Ferron, 2016; Ferron et al., 2020a). Therefore, we asked whether FMRP IF in nociceptor spinal branches localizes to specific axonal microdomains in human spinal cord (**Figure 5**). To do this, we took advantage of the well-established punctate morphology FMRP IF signal in axons (see methods) and the size of primary nociceptor arbors in the dorsal root of human spinal cord (Akins et al., 2017; Christie et al., 2009; Chyung et al., 2018; Gingrich et al., 2018; Monday et al., 2022b). On average, we found that these arbors were roughly 3 μm wide by peripherin staining (cross sectional and short longitudinal axes) (**Figure 5 A-B**). We reasoned that this would permit detection of FMRP puncta (at least 1 μm^2^ signal area) that are present at intra-axonal sites that are spatially associated with either the internal axoplasm (center) or periaxoplasm (plasma membrane) domains (Koenig et al., 2000; Kun et al., 2007; Sotelo-Silveira et al., 2008). To test this, we immunostained for peripherin (to identify internal axoplasm domains) and Nav1.7 (to distinguish periaxoplasm domains) along with FMRP (7G1-1) (Shiers et al., 2021). As shown previously, we found that peripherin IF exhibited cytoplasmic localization and that Nav1.7 IF mostly exhibited plasma membrane localization in dorsal root (Marker colocalization: ~24.7% of Nav1.7; ~25.8% of peripherin) (**Figure 5 insets *i-iii***) (Shiers et al., 2021). We then examined the proportion of FMRP puncta that colocalized with Nav1.7 signal compared to that colocalized with peripherin signal in these arbors from male and female spinal cord (**Table 5**). In male dorsal root, we found approximately 16% of FMRP puncta colocalized with Nav1.7 IF while about 9% colocalized with peripherin IF on average (**Figure 5 A, C**). At high magnification, these puncta often appeared to overlap with both Nav1.7 and peripherin signals (**Figure 5 A *i-iii***). In females, we found about 26% of FMRP puncta were colocalized with Nav1.7 IF and about 15% with peripherin IF on average (**Figure 5 B, C**). These puncta appeared to colocalize with either Nav1.7 or peripherin signal at high magnification (**Figure 5 B *i-iii***). This means about 75% of FMRP puncta in males and about 60% in females did not colocalize with either marker demonstrating that a large proportion do not colocalize with either of these markers. Some of these puncta are likely present in axons that are not of DRG origin like those projected from paravertebral sympathetic ganglia (An et al., 2016; Murata et al., 2003). For detection, our quantification criteria require a perfect 1 μm^2^ overlap or more between FMRP puncta and marker signals omitting puncta that partially overlap with one or both stains (see methods). If this is systematically reducing detected colocalization, then a larger proportion of colocalized FMRP puncta would be detected with a nociceptor receptor stain that is abundant throughout axoplasmic domains, such as TRPV1 (Shiers et al., 2021). This receptor is highly abundant in the plasma membrane and on intracellular vesicles (Bernardini et al., 2004). Nevertheless, these data indicate that FMRP is likely present within both internal axoplasm- and periaxoplasm-associated microdomains in the spinal branches of human nociceptors.

**Figure 5.**
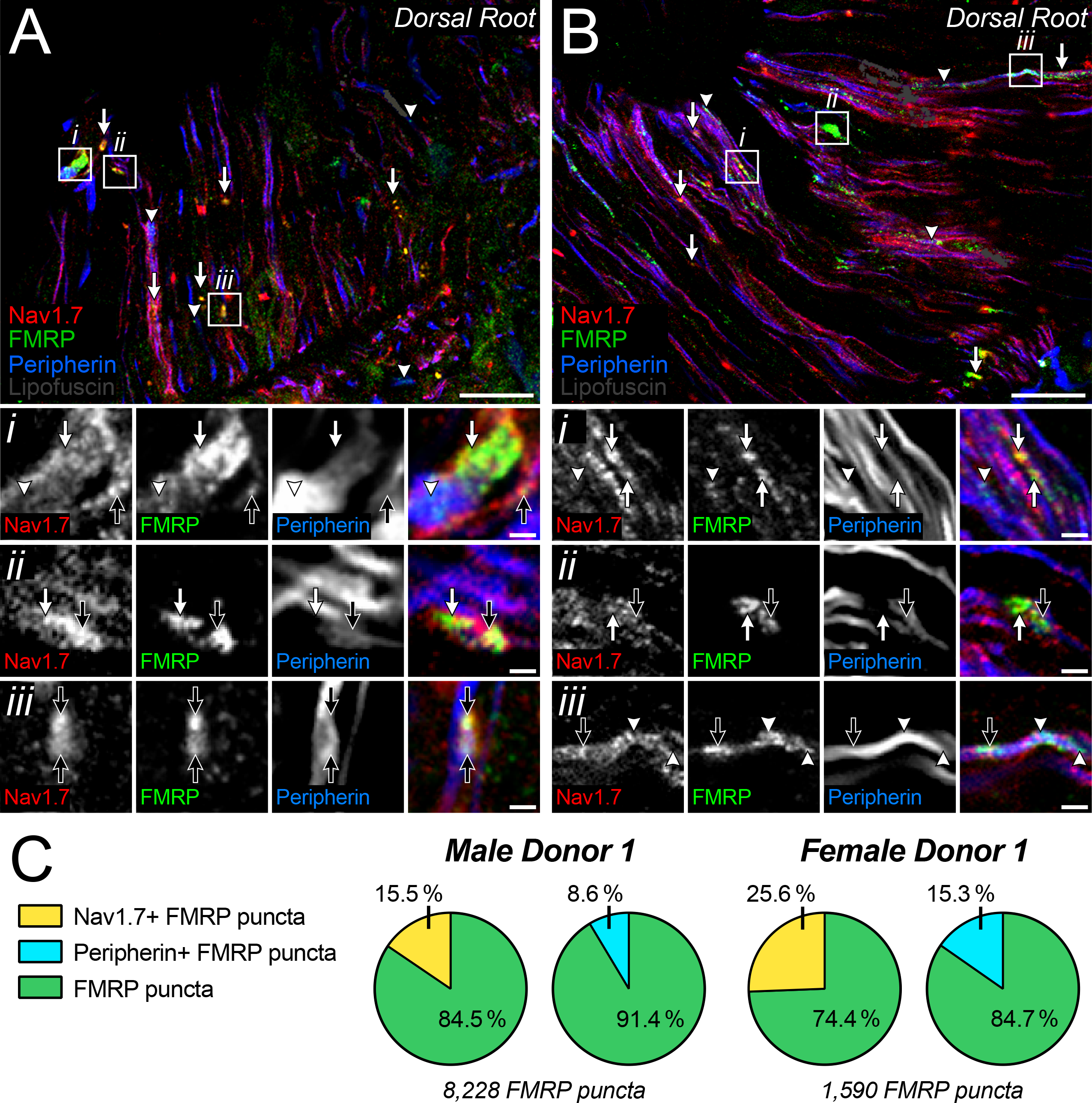
Colocalization of Nav1.7 and/or peripherin with FMRP immunofluorescence in dorsal root. **A-B:** Representative confocal micrographs of immunostaining for Nav1.7 (**Red**), FMRP (**Green**), and peripherin (**Blue**) with lipofuscin (**Grey**) in dorsal root in male (**A**) and female (**B**) human spinal cord. Magnified 40× images representing dorsal root ROIs used to detect FMRP puncta colocalization with Nav1.7 (**Arrows**) and/or peripherin (**Arrowheads**) staining. ***i-iii:*** Further magnified insets (**Boxes**) highlight the sub-axonal organization of these stains. and show varying colocalization amongst all three signals within an arbor (**Black Arrows**). **A:** Colocalized FMRP puncta often overlapped with Nav1.7+ and peripherin+ loci in dorsal root arbors from male spinal cord. **B:** Colocalized FMRP puncta often overlapped with Nav1.7+ or peripherin loci in dorsal root arbors from female spinal cord. **C:** Pie charts showing the average proportion of FMRP puncta in dorsal root per male or female donor that colocalized with Nav1.7 and/or peripherin IF. Scale bars: Magnified Images = 10 μm; Insets = 3 μm.

**Table 4.** Adjustments for potential FMRP puncta colocalization occurring by chance

| Figure | Circuit ROI | Donor | Nociceptor Marker | Total Colocalized FMRP Puncta (#) | Total Adjusted FMRP Puncta (#) | Retention (%) |
| --- | --- | --- | --- | --- | --- | --- |
| 5 | Dorsal Root | Male 2 | Nav1.7 | 13,490 | 6,360 | 47.1 |
|  |  |  | Peripherin | 11,346 | 3,559 | 31.4 |
|  |  | Female 1 | Nav1.7 | 1,569 | 1,221 | 77.8 |
|  |  |  | Peripherin | 1,135 | 733 | 64.6 |
| 6 | Dorsal Root | Male 2 | TRPV1 | 16,502 | 8,455 | 51.2 |
|  |  | Male 3 |  | 11,825 | 7,801 | 66.0 |
|  |  | Female 1 | TRPV1 | 4,919 | 4,297 | 87.4 |
|  |  | Female 2 |  | 5,945 | 3,369 | 56.7 |
|  | Lateral Tracts | Male 2 | TRPV1 | 50,559 | 15,199 | 30.1 |
|  |  | Male 3 |  | 4,842 | 3,632 | 75.0 |
|  |  | Female 1 | TRPV1 | 565 | 146 | 25.8 |
|  |  | Female 2 |  | 9,468 | 5,190 | 54.8 |
|  | Laminae I-II | Male 2 | TRPV1 | 135,550 | 31,181 | 23.0 |
|  |  | Male 3 |  | 348,933 | 162,448 | 46.6 |
|  |  | Female 1 | TRPV1 | 15,147 | 8,428 | 55.6 |
|  |  | Female 2 |  | 140,603 | 41,217 | 29.3 |
| 7 | Dorsal Root | Male 2 | CGRP | 68 | 0 | 0.0 |
|  |  | Male 3 |  | 0 | 0 | 0.0 |
|  |  | Male 4 |  | 1,947 | 1,756 | 90.2 |
|  |  | Female 1 | CGRP | 8,251 | 3,293 | 39.9 |
|  |  | Female 3 |  | 8,415 | 4,642 | 55.2 |
|  |  | Female 4 |  | 8,748 | 5,909 | 67.5 |
|  | Laminae I-II | Male 2 | CGRP | 13,328 | 2,102 | 15.8 |
|  |  | Male 3 |  | 2,446 | 505 | 20.6 |
|  |  | Male 4 |  | 12,673 | 6,826 | 53.9 |
|  |  | Female 1 | CGRP | 33,908 | 16,834 | 49.6 |
|  |  | Female 3 |  | 23,972 | 8,285 | 34.6 |
|  |  | Female 4 |  | 126,996 | 35,098 | 27.6 |

\*Note: The likelihood of FMRP puncta and nociceptor markers colocalizing by chance within an FOV was calculated and then subtracted from the colocalized quantitative data to yield an adjusted proportion of colocalized FMRP puncta per ROI. This was performed using lipofuscin-subtracted puncta numbers from Table 3. The example FMRP puncta numbers given are the overall sums quantified before and after adjustment per donor for each quantitative colocalization analysis.

**Table 5.**
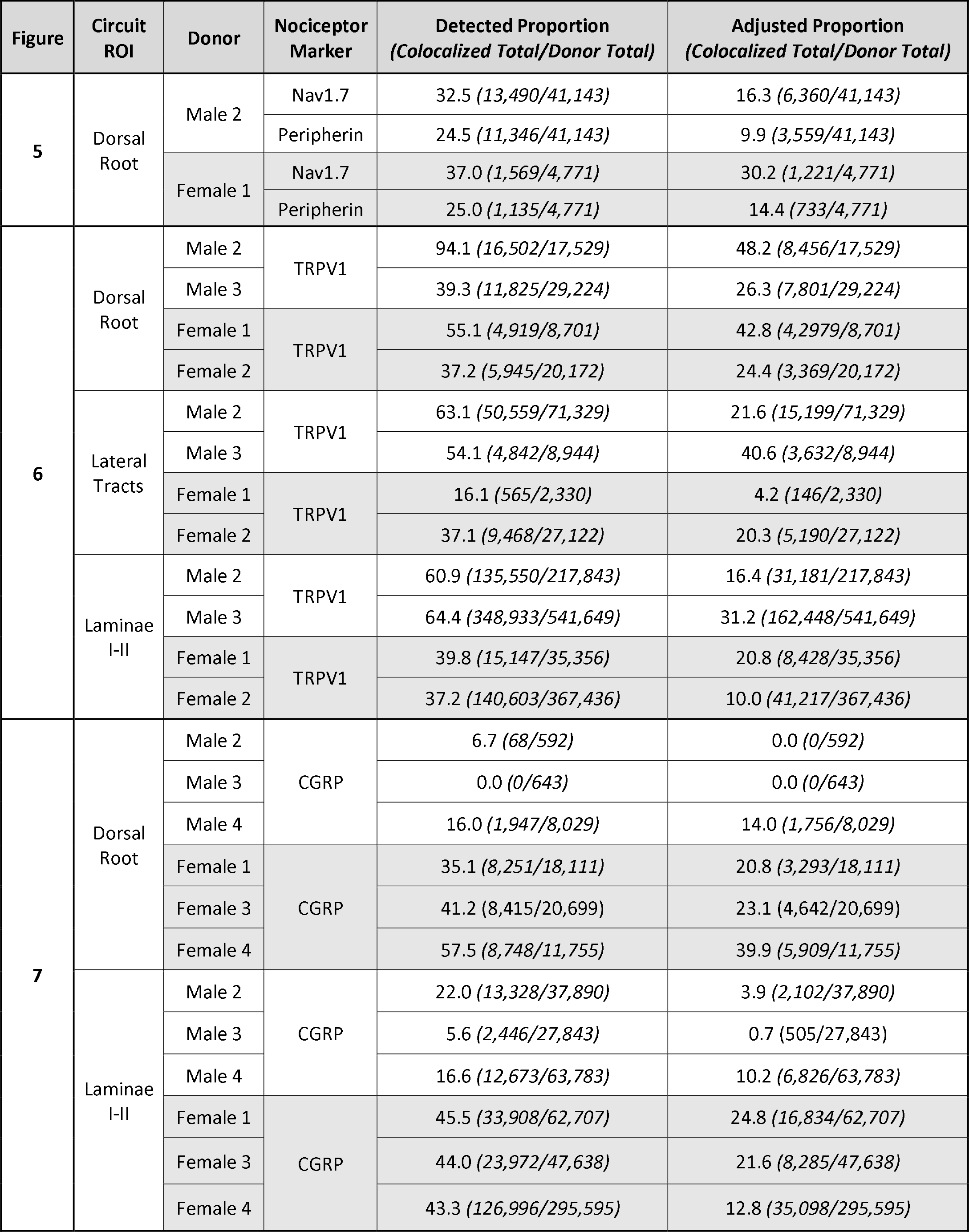
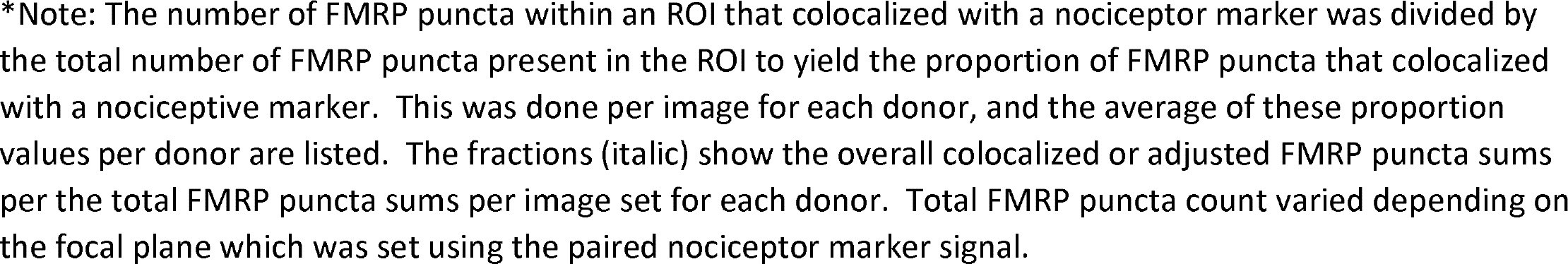
Summarized proportion of FMRP puncta and nociceptive marker colocalization per donor

| Figure | Circuit ROI | Donor | Nociceptor Marker | Detected Proportion<br>(Colocalized Total/Donor Total) | Adjusted Proportion<br>(Colocalized Total/Donor Total) |
| --- | --- | --- | --- | --- | --- |
| 5 | Dorsal Root | Male 2 | Nav1.7 | 32.5 (13,490/41,143) | 16.3 (6,360/41,143) |
|  |  |  | Peripherin | 24.5 (11,346/41,143) | 9.9 (3,559/41,143) |
|  |  | Female 1 | Nav1.7 | 37.0 (1,569/4,771) | 30.2 (1,221/4,771) |
|  |  |  | Peripherin | 25.0 (1,135/4,771) | 14.4 (733/4,771) |
| 6 | Dorsal Root | Male 2 | TRPV1 | 94.1 (16,502/17,529) | 48.2 (8,456/17,529) |
|  |  | Male 3 |  | 39.3 (11,825/29,224) | 26.3 (7,801/29,224) |
|  |  | Female 1 | TRPV1 | 55.1 (4,919/8,701) | 42.8 (4,297/8,701) |
|  |  | Female 2 |  | 37.2 (5,945/20,172) | 24.4 (3,369/20,172) |
|  | Lateral Tracts | Male 2 | TRPV1 | 63.1 (50,559/71,329) | 21.6 (15,199/71,329) |
|  |  | Male 3 |  | 54.1 (4,842/8,944) | 40.6 (3,632/8,944) |
|  |  | Female 1 | TRPV1 | 16.1 (565/2,330) | 4.2 (146/2,330) |
|  |  | Female 2 |  | 37.1 (9,468/27,122) | 20.3 (5,190/27,122) |
|  | Laminae I-II | Male 2 | TRPV1 | 60.9 (135,550/217,843) | 16.4 (31,181/217,843) |
|  |  | Male 3 |  | 64.4 (348,933/541,649) | 31.2 (162,448/541,649) |
|  |  | Female 1 | TRPV1 | 39.8 (15,147/35,356) | 20.8 (8,428/35,356) |
|  |  | Female 2 |  | 37.2 (140,603/367,436) | 10.0 (41,217/367,436) |
| 7 | Dorsal Root | Male 2 | CGRP | 6.7 (68/592) | 0.0 (0/592) |
|  |  | Male 3 |  | 0.0 (0/643) | 0.0 (0/643) |
|  |  | Male 4 |  | 16.0 (1,947/8,029) | 14.0 (1,756/8,029) |
|  |  | Female 1 | CGRP | 35.1 (8,251/18,111) | 20.8 (3,293/18,111) |
|  |  | Female 3 |  | 41.2 (8,415/20,699) | 23.1 (4,642/20,699) |
|  |  | Female 4 |  | 57.5 (8,748/11,755) | 39.9 (5,909/11,755) |
|  | Laminae I-II | Male 2 | CGRP | 22.0 (13,328/37,890) | 3.9 (2,102/37,890) |
|  |  | Male 3 |  | 5.6 (2,446/27,843) | 0.7 (505/27,843) |
|  |  | Male 4 |  | 16.6 (12,673/63,783) | 10.2 (6,826/63,783) |
|  |  | Female 1 | CGRP | 45.5 (33,908/62,707) | 24.8 (16,834/62,707) |
|  |  | Female 3 |  | 44.0 (23,972/47,638) | 21.6 (8,285/47,638) |
|  |  | Female 4 |  | 43.3 (126,996/295,595) | 12.8 (35,098/295,595) |

\*Note: The number of FMRP puncta within an ROI that colocalized with a nociceptor marker was divided by the total number of FMRP puncta present in the ROI to yield the proportion of FMRP puncta that colocalized with a nociceptive marker. This was done per image for each donor, and the average of these proportion values per donor are listed. The fractions (*italic*) show the overall colocalized or adjusted FMRP puncta sums per the total FMRP puncta sums per image set for each donor. Total FMRP puncta count varied depending on the focal plane which was set using the paired nociceptor marker signal.

### 3.5 FMRP localizes to TRPV1+ and CGRP+ axons in the dorsal horn

Nav1.7 and peripherin localize to human nociceptors but are also expressed by non-nociceptive neurons in humans while TRPV1 and CGRP are more specific to nociceptor populations (Shiers et al., 2021; Tavares-Ferreira, Shiers, et al., 2022). To identify the relative proportion of FMRP that is present in nociceptor axons, we analyzed colocalization between TRPV1 IF and FMRP puncta in dorsal root, lateral white matter tracts, and laminae I-II (**Figure 6; Table 5**). To detect TRPV1+ axons, we used an antibody previously characterized to detect TRPV1 receptors in human DRG and spinal cord (**Table 2**) (Shiers et al., 2021). We found approximately 35% of the FMRP puncta present in dorsal root colocalized with TRPV1+ axons (**Figure 6 A, D**). In lateral white matter tracts, we found 22% colocalization (**Figure 6 B, D**). In laminae I-II, we found roughly 20% of the total FMRP puncta detected colocalized with TRPV1+ fibers (**Figure 6 C, D**). These data show that a substantial fraction of FMRP present in these regions localizes to segments of nociceptor axons that have TRPV1 protein and indicate that FMRP is present throughout the nociceptor axoplasm.

**Figure 6.**
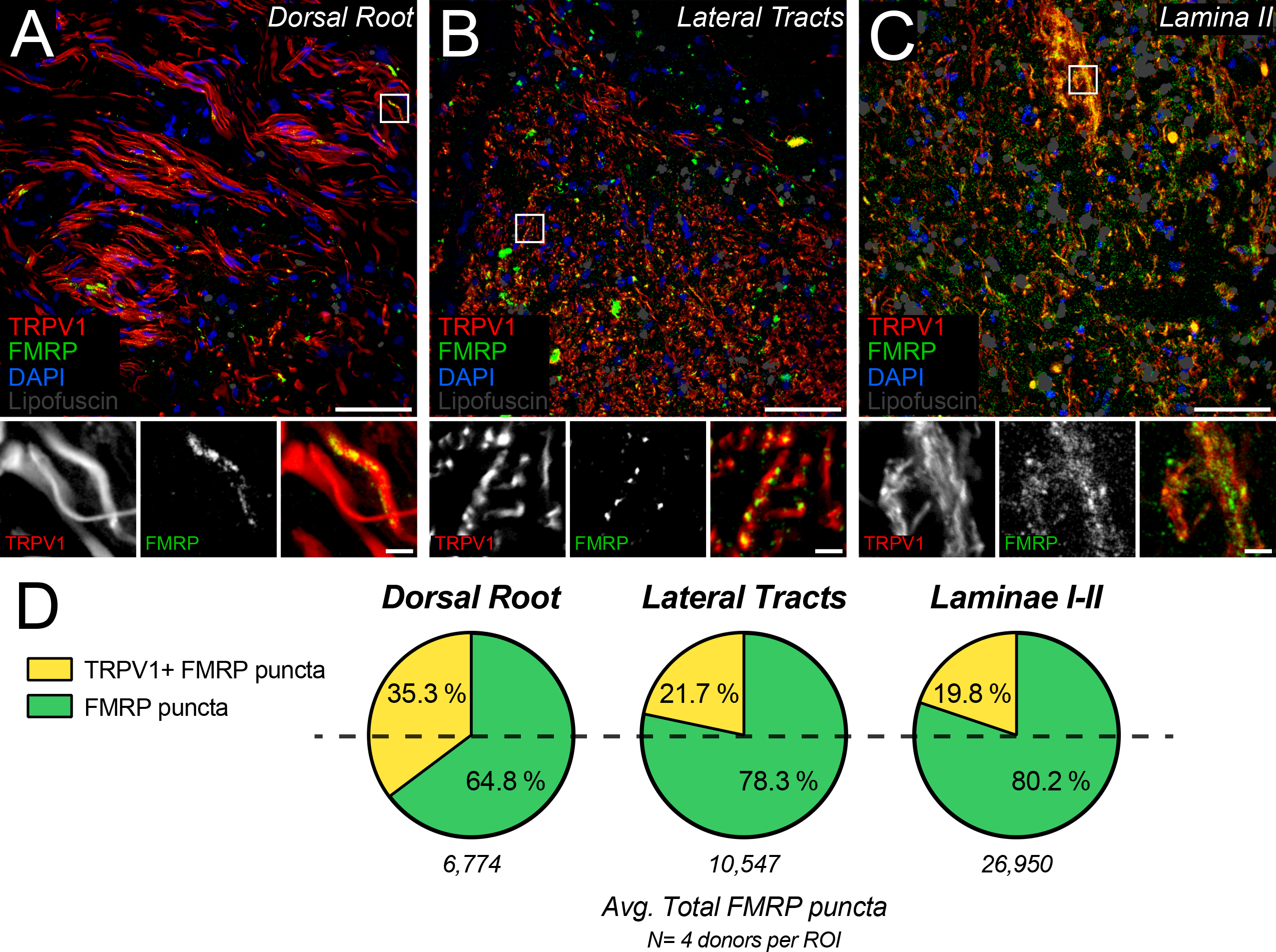
Colocalization of TRPV1 and FMRP immunofluorescence in human nociceptive circuits. **A-C**: Representative confocal micrographs of human nociceptive circuitry as identified by TRPV1 staining (**Red**) with FMRP IF (**Green**), DAPI (**Blue**), and identified lipofuscin (**Grey**) in human spinal cord. Magnified 40x images represent nociceptive circuit ROIs used to detect the proportions of FMRP puncta present that colocalized with TRPV1+ fibers. Associated insets (**Boxes**) are further magnified to highlight the sub-axonal morphology of this colocalization. **A:** Image of dorsal root shows FMRP puncta frequently colocalized with TRPV1+ axons. Glial nuclei rarely colocalized with FMRP. **B:** Image of lateral tracts shows FMRP puncta often colocalized with these TRPV1+ arbors. **C:** Image of lamina I-II neuropil showing notable colocalization between FMRP puncta and TRPV1+ fibers in these synaptic fields. **D:** Pie charts showing the average proportion of FMRP puncta per nociceptive circuit ROI the did or did not colocalize with peripherin IF. Scale bars: Magnified Images = 50 μm; Insets = 5 μm.

We then asked whether FMRP colocalizes with nociceptor axons and terminals containing the neuropeptide, calcitonin gene-related peptide (CGRP). Unlike in mice, the majority of nociceptor somata in human DRG contain CGRP, and this population overlaps almost entirely with TRPV1 which is expressed by virtually all human nociceptors (Shiers et al., 2021). However, at the subcellular level in the substantia gelatinosa, CGRP IF is detected at distinct presynaptic loci in nociceptor branches (Shiers et al., 2021). This potentially arises from the spatial segregation of neuropeptide-containing vesicles to different sites of these terminals, such as presynaptic glomeruli (McNeill et al., 1988; X. Zhang et al., 1995). Consistent with prior findings, we detected fewer CGRP+ axons in dorsal roots when compared to TRPV1+ axons, but like TRPV1 IF, found CGRP signal throughout the entirety of laminae I-II (**Figure 7 A-D**) (Shiers et al., 2021). Interestingly, our analysis revealed a stark difference in FMRP IF colocalization with these sites between male and female samples (**Figure 7 E-F; Table 5**). In spinal cord sections recovered from male donors, we found little colocalization in dorsal root (about 7.0%) and only 5.2% in laminae I-II (**Figure 7 A-B, E**). However, in female donors, we readily detected about 27.9% colocalization with the few CGRP+ axons present in dorsal root (**Figure 7 C, E**). In laminae I-II, we found about 18.9% colocalization on average (**Figure 7 D, E**). This difference between male and female donors was significantly different by t-test with *p*-values of 0.0375 for dorsal root and 0.0310 for laminae I-II (**Figure 7 F**). These data indicate that synaptic regulation of FMRP and its engagement in CGRP-containing spinal terminals are likely prominent mechanisms in human nociceptor axons with potential female-selective actions in nociceptive plasticity.

**Figure 7.**
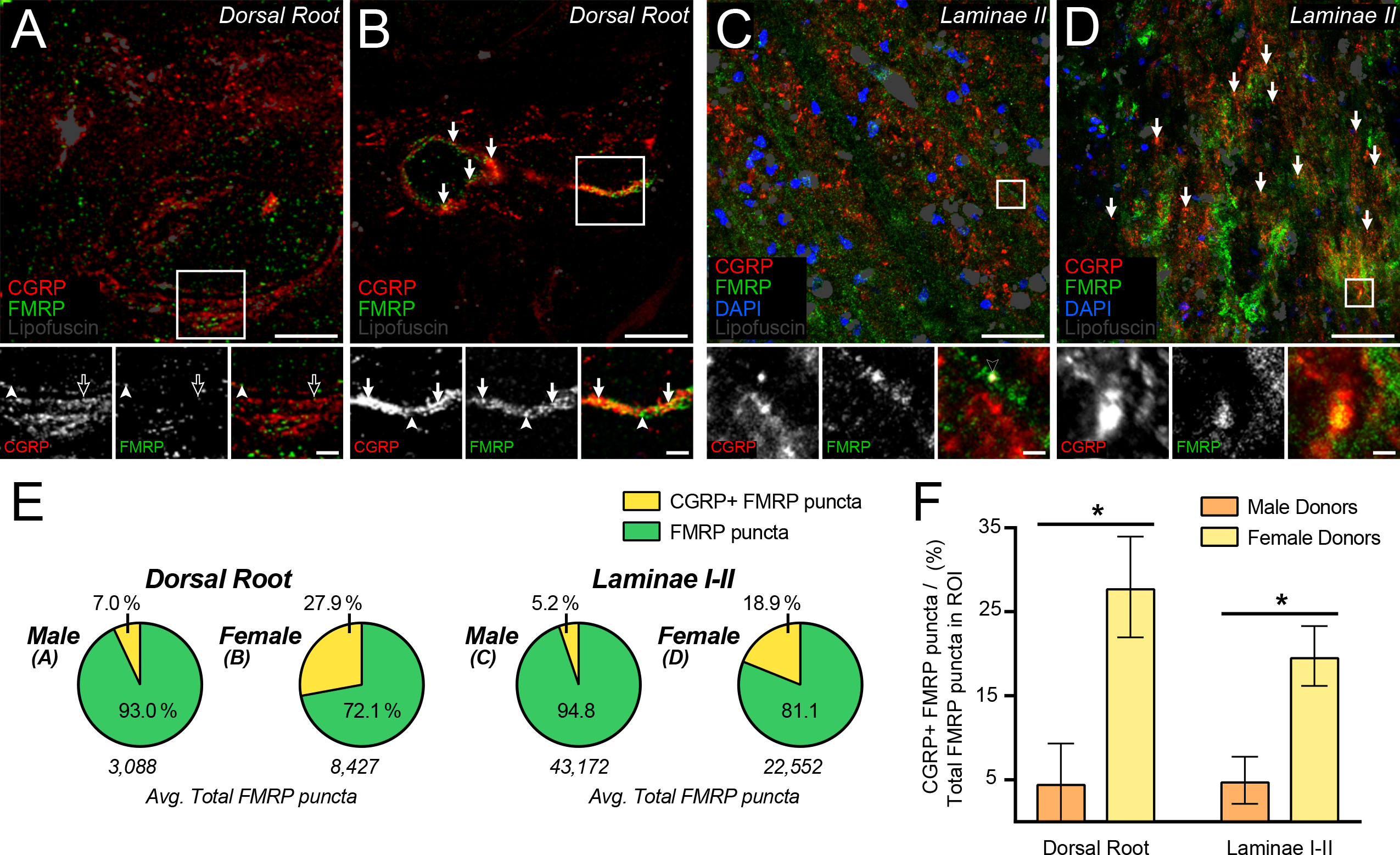
Colocalization of CGRP and FMRP immunofluorescence in dorsal root and laminae I-II. **A-D:** Representative confocal micrographs of CGRP (**Red**) and FMRP (**Green**) immunofluorescence with lipofuscin (**Grey**) in dorsal root (**A-B**) and laminae I-II (**C-D**) in male and female human spinal cord. Magnified 40× images representing ROIs used to detect colocalization between CGRP and FMRP. Colocalization was notably rare in tissues from male donors but was notably frequent in tissues from female donors (**Arrows**). Associated insets (**Boxes**) are further magnified highlighting the sub-axonal organization of these stains. **A:** Image from a male donor showing that almost all FMRP puncta are spatially separated from CGRP+ loci. Few puncta outside our detection parameters rarely colocalize with CGRP staining (**Black Arrow**). **B:** Image from a female donor showing FMRP puncta frequently colocalized with CGRP+ loci. Magnified 40x images of laminae I-II were counterstained with DAPI (**Blue**) to show neuropil and somatic organization. **C:** Image from a male donor showing that almost all FMRP puncta are spatially separated from CGRP+ loci as in dorsal root. >Potential colocalized puncta were often identified as lipofuscin (**Black Arrowhead**). **D:** Image from a female donor showing frequent colocalization between CGRP and FMRP puncta. **E:** Pie charts showing the average proportion of FMRP puncta per ROI the did or did not colocalize with CGRP+ loci. **F:** Comparisons showed that colocalization between FMRP puncta and CGRP+ loci was significantly increased in females when compared to males for both dorsal root (p= 0.0375) and laminae I-II (p= 0.0310) analyses. Values expressed as mean +/− SEM. T-test: \**p* < 0.05 (N = 3 per group). Dorsal root images = 15 μm; Laminae I-II images = 50 μm; All insets = 5 μm.

## 4 Discussion

We have demonstrated that FMRP, the critical regulator of local translation and synaptic plasticity, is expressed in human nociceptor axons of the dorsal horn. Our analyses revealed it is present in human DRG and spinal neurons. In spinal neuropil, FMRP immunoreactivity is most abundant in substantia gelatinosa. FMRP puncta are found in nociceptor axons within dorsal root, lateral white matter tracts, and laminae I-II but not in dorsal funiculus arbors. Unexpectedly, these puncta colocalized with CGRP+ loci in nociceptor axons from female but not male donors, indicating its spinal presynaptic localization is likely sex dimorphic. Overall, our findings are consistent with FMRP regulating axonal translation and presynaptic plasticity in these branches.

FMRP is present in most human DRG neurons. This is consistent with its mRNA and protein expression in human and rodent DRG neurons, respectively (Price et al., 2006; Tavares-Ferreira, Shiers, et al., 2022). Studies in FXS and premutation patients suggest that FMRP plays a bi-directional role in pain (Au et al., 2013; Hagerman et al., 2007; Johnson et al., 2022; Leehey et al., 2011; Lozano et al., 2016; Salcedo-Arellano et al., 2020; Symons et al., 2003). In rodents, FMRP regulates DRG neuron excitability and long-term plasticity including in chronic pain models (Asiedu et al., 2011; P.-Y. Deng et al., 2021; Ferron et al., 2014; Ferron, 2016; Ferron et al., 2020a; Price et al., 2007). *FMR1* null mice exhibit decreased acute nociceptive sensitization, reduced inflammation-induced hyperalgesia, and delayed SNI-induced mechanical allodynia (Asiedu et al., 2011; Price et al., 2007). This arises in part from decoupling of activity-dependent programs from extracellular signals in DRG afferents (Price et al., 2007). Our findings suggest that FMRP is well-positioned to play a similar role in human nociceptors.

FMRP is also present in human spinal neurons and neuropil, being most evidently abundant in laminae I-II synaptic fields. This is consistent with its role in rodent dorsal horn in inflammatory and neuropathic pain (Price et al., 2007; Ramírez-López et al., 2021; Y. Yang et al., 2021). In CNS circuits, its best-studied function is coupling local translation to receptor activation *(e.g., mGluR1/5; NMDAR)* at postsynaptic sites to rapidly modify long-term synaptic function in brain (Antar & Bassell, 2003; Bagni & Zukin, 2019; Bear et al., 2004; Kute et al., 2019; Paul et al., 2019; Routh et al., 2013; Zalfa et al., 2003). This pairs synaptic activity with local synthesis of receptor/channel subunits *(e.g., AMPAR; Kv4.2)* and internalization machineries *(e.g., Arc; SYNGAP1; SAPAP3)* to modulate dendritic channel and receptor surface expression (Antar et al., 2004; Bagni & Zukin, 2019; Bear et al., 2004; Niere et al., 2012; Paul et al., 2019; Routh et al., 2013; Wang et al., 2008, 2010). These receptors/channels play critical roles in chronic pain-associated spinal circuit activity, and these mRNAs are highly expressed in human nociceptors (Harte et al., 2018; Iyengar et al., 2017; Latremoliere & Woolf, 2009; Ray et al., 2022; Sluka & Clauw, 2016; Tavares-Ferreira, Shiers, et al., 2022).

FMRP is present along the extent of centrally-projecting human nociceptor axons targeting substantia gelatinosa. In rodents, these branches contain translation regulators including FMRP, polyribosomes, and mRNA (Koenig et al., 2000; Paige et al., 2020; Patil et al., 2019; Price et al., 2006). FMRP directly associates with polyribosomes in ribonucleoprotein particles (RNPs) and binds over 1000 different mRNA species in brain (Ascano et al., 2012; Bagni & Zukin, 2019; V. Brown et al., 2001; Darnell et al., 2011; Darnell & Klann, 2013; Hale et al., 2021; Sawicka et al., 2019). Its target mRNAs are necessary for long-term plasticity and are predominately synaptic with critical functions in neurotransmission (Bagni & Zukin, 2019; Chyung et al., 2018; Darnell et al., 2011; Hale et al., 2021; Monday et al., 2022b; Sawicka et al., 2019). Evidence in brain circuits shows that axonal translation occurs near boutons and that FMRP regulates this process to modify long-term presynaptic function (Christie et al., 2009; Hafner et al., 2019; Monday et al., 2022b; Scarnati et al., 2018). In hippocampus, recent findings demonstrate that axonal FMRP-RNPs couple circuit activity to axonal protein synthesis and long-term presynaptic remodeling required for LTP (Monday et al., 2022b). In rodent and human brain circuits, FMRP-RNPs contain mRNAs critical for bouton formation and synaptic vesicle dynamics that are also highly expressed in subsets of human nociceptors (Akins et al., 2017; Bamji et al., 2003, 2006; Chyung et al., 2018; Ray et al., 2022; Sun et al., 2009; Sun & Bamji, 2011; Tavares-Ferreira, Shiers, et al., 2022; Taylor et al., 2013). Long-term nociceptor plasticity also requires axonal translation, a well-characterized mechanism in their peripheral arbors that reorganizes the local proteome and enhances local signaling (L. F. Ferrari et al., 2015; Khoutorsky & Price, 2018; Price & Géranton, 2009; Reichling & Levine, 2009). Since FMRP is present in human nociceptor fibers in substantia gelatinosa, it may associate with RNPs and mRNAs critical for presynaptic function near their boutons as shown in other CNS circuits. This would couple peripheral and spinal signaling to axonal translation and long-term enhancements in neurotransmission from nociceptors to nociceptive circuits.

In dorsal root, FMRP puncta were present in plasma membrane-associated microdomains. This is consistent with its binding and modulation of ion channels in the axonal plasma membrane (Brager & Johnston, 2014; Contractor, 2013; P.-Y. Deng et al., 2013, 2019, 2021; Ferron, 2016; Ferron et al., 2020b). In DRG neuron terminals, it directly binds Cav2.2 to regulate synaptic vesicle exocytosis (Ferron, 2016; Ferron et al., 2014, 2020b). FMRP may have similar functions in human nociceptor fibers innervating substantia gelatinosa. Since it binds channels and RNA with distinct motifs, this has led to a growing model in CNS circuits whereby FMRP regulates the spatial pairing of axonal RNPs with nearby channel signaling by binding both simultaneously (Akins et al., 2017; Bagni & Zukin, 2019; P.-Y. Deng & Klyachko, 2021; Ferron, 2016; Korsak et al., 2016; Monday et al., 2022b). Activity-driven transport or mobility of channels or channel pools therefore likely mediates synapse-specific targeting of FMRP and associated polyribosomes, a well-documented observation in dendrites (Auerbach & Bear, 2010; Bear et al., 2004; Bhakar et al., 2012; F. Ferrari et al., 2007). In line with this, FMRP puncta were also present in axoplasmic microdomains of these axons in dorsal root and in fibers in lateral tracts and laminae I-II. This suggests that FMRP in these axonal compartments is being targeted to synapses along with its mRNA cargoes potentially in response to synaptic activity.

FMRP puncta colocalized with CGRP+ loci in the spinal branches of primary nociceptors in female but not male donor spinal cord. Maladaptive changes in CGRP signaling are emerging as a feature of many chronic pain phenotypes specifically in females (Avona et al., 2019, 2021; Casale et al., 2021; Iyengar et al., 2017; KORUCU et al., 2020; Paige et al., 2022; Siracusa et al., 2021; Sluka & Clauw, 2016; Tavares-Ferreira, Ray, et al., 2022). In female rodents, spinal CGRP signaling plays an important role in multiple chronic pain models by regulating excitatory/inhibitory balance in intrinsic spinal cord neurons (Paige et al., 2022). Shifts in CNS CGRP release are thought to arise at least in part from sex differences in spinal synaptic loci of NMDA receptor (NMDAR) signaling (Iyengar et al., 2017; Sluka & Clauw, 2016; Temi et al., 2021). NMDAR are major drivers of experience-dependent FMRP accumulation in sensory circuits that engage FMRP-regulated translation (Gabel, 2004; Kute et al., 2019; Martin et al., 2016; Paul et al., 2019; Routh et al., 2013; Thomazeau et al., 2021; Todd et al., 2003; Yau et al., 2016). At the synapse, NMDAR activity modifies FMRP phosphorylation altering its association with local RNPs and enhancing local protein synthesis (Kute et al., 2019; Paul et al., 2019; Routh et al., 2013; Thomazeau et al., 2021). On boutons, NMDARs drive synapse-specific localization of RNA granules (Wong et al., 2022). NMDARs are also present on the spinal terminals of nociceptors and can potentiate their spinal inputs (M. Deng et al., 2019). In females, our data indicate that an increased amount of FMRP is positioned to be engaged by NMDAR signaling at CGRP-expressing terminals and may regulate long-term synaptic release properties of these boutons, including for CGRP itself. Overall, this suggests that FMRP differentially regulates spinal presynaptic plasticity of human nociceptor axons with a sex dimorphic target being CGRP+ terminals in females.

There are several limitations to this study. First, we acknowledge that the human tissue sample size is small per male and female donor group. We designed our IF studies in human spinal cord from 4 donors (2 male and 2 female) to characterize FMRP expression in spinal nociceptor axons and unexpectedly found a sex difference in FMRP and CGRP colocalization. No previous sex differences in FMRP function or localization have been documented. Second, we did not validate our colocalization findings with electron or super resolution microscopy. We designed these experiments to identify whether FMRP is present in these axons by empirically determining the proportion of FMRP puncta in dorsal horn regions that colocalized with nociceptor markers. Third, we did not verify our antibody specificity in FXS DRG or spinal cord due to limitations in tissue availability. We thoroughly compared the three major monoclonal FMRP antibodies using 2F5-1 as a positive control since it was previously used to detect FMRP in human brain tissues (Akins et al., 2017). Analyses with 7G1-1 were restricted to anatomical sites the showed consistent IF across all three antibodies. Fourth, we did not stain for mRNA or rRNA when identifying FMRP puncta. Future studies will assess this and the regulation of FMRP association. Finally, we have not performed functional experiments, but this is a limitation of working with human tissues from organ donors.

Chronic pain symptoms still severely impair daily life for millions of neuropathic pain patients around the world (Dahlhamer, 2018; Yousuf et al., 2021). Notably, severity is often magnified in women when compared to men (Fillingim et al., 2009; Mogil, 2020; Traub & Ji, 2013). Primary nociceptor activity is required for the generation of most pain symptoms (Haroutounian et al., 2014, 2018; Khoutorsky & Price, 2018; Price & Géranton, 2009; Price & Gold, 2018; Siracusa et al., 2021; Yousuf et al., 2021). Long-term shifts in the function of these neurons requires translation (Fillingim et al., 2009; Mogil, 2020; Traub & Ji, 2013). FMRP is a critical regulator of local translation and synaptic plasticity in humans associated with pain pathophysiology in FXS and premutation patients (Asiedu et al., 2011; Bagni & Zukin, 2019; Darnell & Klann, 2013; Johnson et al., 2022; Mei et al., 2020; Price et al., 2007; Ramírez-López et al., 2021; L. Yang et al., 2020; Y. Yang et al., 2021). Taken with our findings, FMRP is likely a critical regulator of spinal presynaptic plasticity in human nociceptor axons, and notably, is positioned to regulate the sex dimorphic actions of spinal CGRP signaling in nociceptive plasticity and chronic pain.

## Acknowledgments

This work was supported by NIH grants NS065926, NS102161 and NS111929. The authors are grateful to the organ donors and their families for the gift of life and research provided by their organ donation.

